# Environmental selection overrides host selection in a horizontally acquired microbiota

**DOI:** 10.1101/2023.03.22.533819

**Authors:** Nancy Obeng, Hinrich Schulenburg

## Abstract

Horizontally acquired symbionts need to succeed both within hosts and the free-living environment. Microbes might invest differentially in these habitats, thus shaping fitness within host-associated microbiota communities. In this study, we investigated how host and environmental selection affect microbiota composition in a two-member model community of *Pseudomonas lurida* MYb11 and *Ochrobactrum vermis* MYb71 from the natural microbiota of *Caenorhabditis elegans*. Fitness characterizations in the host and non-host environment revealed distinct ecological strategies: while MYb11 dominated free-living by rapidly growing, it was quickly outcompeted within worms by MYb71. Using mathematical modeling and experimental evolution, we assessed how these distinct strategies affect microbiota composition over time. We found that MYb11 enriches in the host via its advantage in the environment and additionally adapts to the host. This community shift was mirrored in host fitness. Overall, this highlights the importance of the symbiont pool and shows that environmental selection can overwhelm host adaptation.

## Introduction

Animals can horizontally acquire individual symbionts or whole microbiota communities from the environment. In host-association, these microbes provide nutrients, influence host physiology, development and reproduction and thereby ultimately shape organismal fitness^1, 2^. This requires microbes to colonize novel hosts, maintain the association and eventually transmit to the next. For successful transmission, they additionally need to proliferate or at least survive in the free-living environment. In such a biphasic life cycle, microbes are thus challenged with selection by two distinct compartments: the host and the environment^3^. While considerable attention has focused on host-association strategies of microbes (e.g. ^3–5^), it remains unclear how environmental and host selection balance and determine microbiota composition and function.

Selection and diversification are two key processes that shape communities. While selection acts on deterministic fitness differences between individuals from different species, diversification describes the emergence of new genetic variation^6–8^. Naturally, these processes also occur in microbiota communities, where short-term ecological and long-term evolutionary dynamics often overlap^7, 9^. When microbiotas are acquired horizontally, microbes are exposed to selection both in the host and the environment^3^. In the short term, species sorting should drive microbiota composition as bacteria specializing on either habitat may face costs in the other. In the long term, i.e. over many generations, adaptive change of individual lineages can additionally change microbiota composition. However, relatively little is known about how host and environmental selection balance during microbiota assembly and evolution. Recent theoretical work suggests that a single species evolving in a biphasic life cycle should either (i) invest in increased growth in the environment or (ii) decrease migration from host, i.e. stay, to optimize fitness^10^. However, it is unclear how strategies of co-existing microbes affect an evolving community. Experimentally, the complexity of most microbiotas makes it difficult to assess the relative influence of host and environmental selection as biotic interactions between microbiota members influence fitness, yet become non-linear with increasing community complexity^11^.

In this study, we examined a simple two-member microbiota model to assess how fitness investments in the free-living vs. host-associated phase shape microbiota composition along the biphasic life cycle on ecological and evolutionary timescales. We used a well-characterized pair of bacteria from the natural microbiota of the nematode *Caenorhabditis elegans*^12–14^: *Pseudomonas lurida* MYb11, which can provide immune-protection against pathogens but only occasionally associates with *C. elegans*^12, 15–18^, and *Ochrobactrum vermis* MYb71, which efficiently colonizes the nematode^12, 18–21^. Originally co-isolated from the natural worm strain *C. elegans* MY316 (*Ce*_MY316)^12^, they can co-exist within worms and on agar^12–14^. In the current study, we used mono- and co-culture experiments to quantify relative bacterial fitness along a biphasic life cycle, in the host *Ce*_MY316 and free-living on agar. We further tested whether the combination of relative fitness in the host and environment dictates changes in community composition over repetitions of the life cycle using a mathematical model and experimental evolution. Finally, we evaluated how changes in the two-member community influence host fitness. Together, this provides a new perspective on the drivers and implications of microbiota change.

## Results

### Context determeading ines interspecies interactions in a two-member microbiota

To assess bacterial fitness of the microbiota isolates *P. lurida* MYb11 and *O. vermis* MYb71 in the host and the free-living environment (Fig. 1A), we quantified colony forming units of both species in mono- and co-association in the two habitats. In the free-living agar environment, they both showed significantly greater population sizes after three days growth in mono-culture than together (Fig. 1B, Supplementary Table S1), indicating competition. We selected this time point as a reference for free-living fitness, as it reflects the duration of the free-living phase in our evolution experiment below. As communities plateaued in size after extended incubation (seven days), this growth cost disappeared (Extended Data Fig. 1, Supplementary Table S1). As expected based on their co-isolation from the natural worm isolate *Ce*_MY316, both species colonized worms alone and in combination. Within worms, however, competition was attenuated, as they reached comparable abundances in mono-as well as in co-colonization (Fig. 1C, Supplementary Table S2). As habitat appeared to shape bacterial fitness and interactions, we continued to map bacterial fitness along the biphasic life cycle in greater detail.

**Figure 1:**
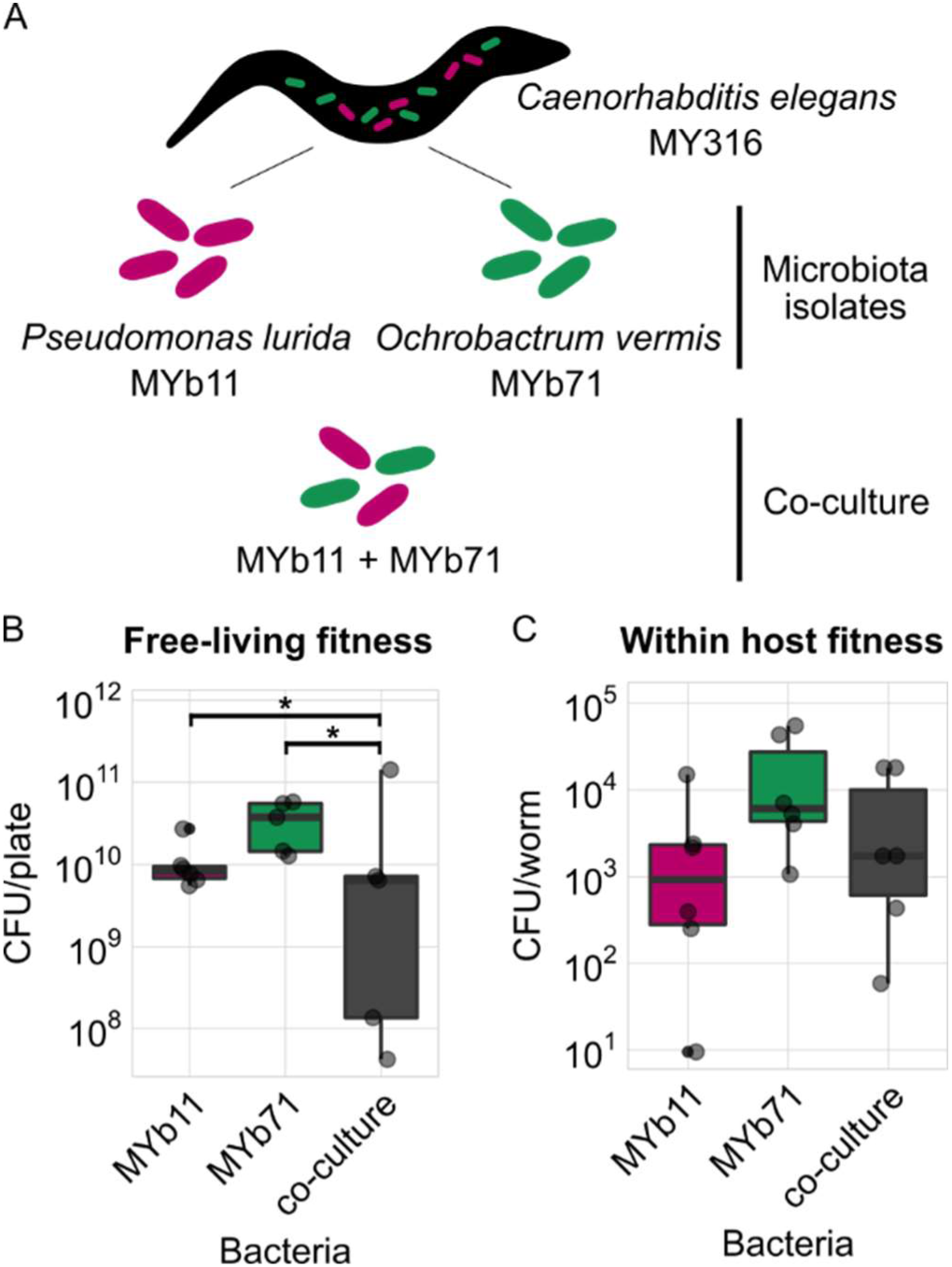
Microbiota members *Pseudomonas lurida* MYb11 and *Ochrobactrum vermis* MYb71 compete in the free-living environment, but not within the host. **A** The model microbiota community used in this study consists MYb11 and MYb71, two key members of the natural microbiota of *Caenorhabditis elegans*. They are investigated in their natural host, *C. elegans* strain MY316 (*Ce*_MY316), in mono- and in co-culture. **B** Free-living fitness of bacteria was measured as colony forming units (CFUs) that grew on nematode growth agar within 72h of incubation at 20°C (equivalent to selection time point of the later evolution experiment, see Fig. 3). **C** Within host fitness was measured as CFU/worm collected from L4 stage worms raised on the respective bacteria. **B – C** Replicates are shown as individual data points (5 ≤ n ≤ 6). Box plots show median (center line), upper and lower quartiles (box limits) and the interquartile range (whiskers). Differences between mono- and co-culture were detected using Dunnett tests (* = P < 0.05).

### Microbiota members have distinct fitness strategies along the biphasic life cycle

In a biphasic life cycle, symbionts need to enter hosts, establish and persist in colonization, and release back into environment where they may grow (Fig. 2). To compare fitness strategies within our two-member community, we performed competition experiments along the life cycle. Species abundance (in colony forming units, CFUs) was determined by selective plating and used as a proxy for relative fitness (as in^13, 22, 23^ for example). In the early phase of colonization, i.e. after 1.5 hours of worm exposure, similar numbers of MYb11 and MYb71 were found in L4 stage worms, although MYb11 dominated often (Fig. 2A; Supplementary Table S3). Bacterial communities that had established in L4 larvae had a roughly even species composition, although MYb71 dominated in more samples than MYb11 (Fig. 2B; Supplementary Table S3).

**Figure 2:**
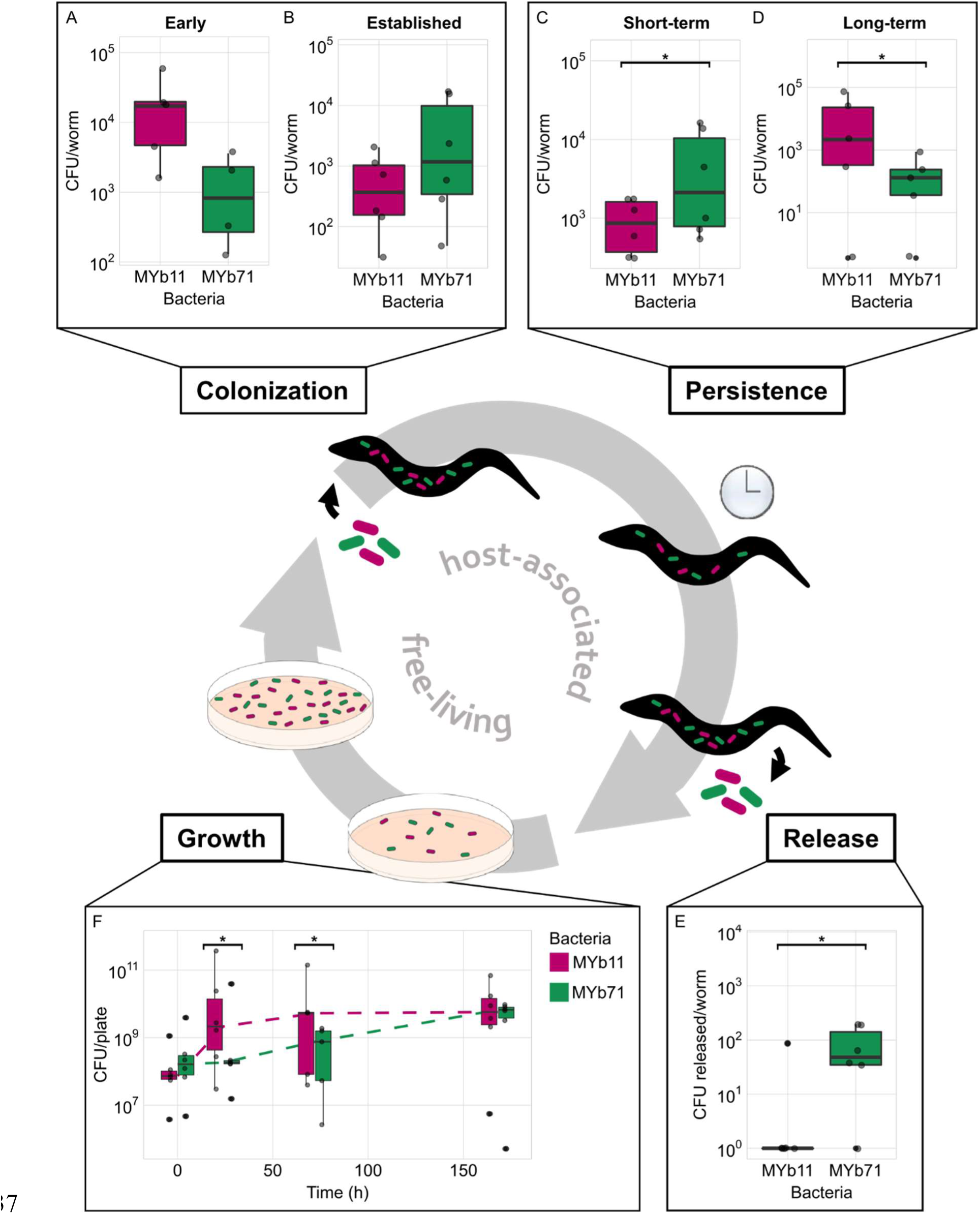
Microbiota members MYb11 and MYb71 have distinct fitness strengths along the biphasic symbiotic life cycle. Bacterial fitness within co-cultures was measured in host-association (CFU per *Ce_*MY316 as a host) or free-living (CFU per population on nematode growth agar). Host-associated life stages include **A** early colonization of L4 stage worms (within 1.5h exposure), **B** established colonization (worms that developed on respective bacteria since L1 stage), **C** short-term and **D** long-term persistence in worms (maintenance after 1h in M9 buffer and 24h on empty nematode growth agar, respectively) and **E** release from L4 stage worms into buffer within 1h. **F** Free-living fitness, i.e. bacterial abundance reached via growth, in the co-culture was assessed over seven days (sampled at 0h, 24h, 72h and 168h). Dashed lines connect median CFU/plate-values between time points. **A – F** Replicates are shown as individual data points (4 ≤ n ≤ 6). Box plots show median (center line), upper and lower quartiles (box limits) and the interquartile range (whiskers). Differences between the bacterial species’ abundances were detected using paired t-tests (* = P < 0.05).

As both early and established colonization were measured in worms that had been directly exposed to lawns of environmental bacteria before the quantification, we further assessed persistence in worms. Here we distinguish between short-term persistence, so bacteria that are not immediately excreted or digested within one hour of worms being away from the environmental symbiont pool, and long-term persistence, so bacteria that survive and are maintained in worms for extended periods of time in the absence of the environmental symbiont pool (24 hours on empty plates). In the short-term, MYb71 significantly enriched in the communities, suggesting that it is better adapted to the worm than MYb11 (Fig. 2C; Supplementary Table S3). Quantifying the CFUs that exited worms in the meantime, we also observed a greater abundance of MYb71 in the collected buffer (Fig. 2E, Supplementary Table S3). This suggests that MYb71 accumulates in worms, yet is efficiently released from the host. Under more extreme conditions of persistence, when *Ce*_MY316 are maintained on empty NGM for 24 hours and are likely starving, MYb11 persists significantly better than MYb71, whose abundance dropped by about one order of magnitude (Fig. 2D; Supplementary Table S3).

In the free-living environment, too, our two focal species showed distinct population dynamics. After 24 and 72 hours of growth, respectively, MYb11 was significantly enriched in the plate environment (Fig. 2F; Supplementary Table S3). This revealed that MYb11 is the dominant competitor in the plate environment during early and mid-term growth (24h and 72h, respectively), where both species suffered a fitness cost when growing together (Fig. 1B). In line with this, MYb11 was also significantly more motile, indicating also greater capacity to spatially access the agar environment (Extended Data Fig. 2; Supplementary Table S4). After a long period of seven days of growth, however, both species reached similar abundances (Fig. 2F; Supplementary Table S3), possibly the carrying capacities. Together, this indicates that a rapid onset of growth provides a competitive advantage to MYb11 in the plate environment, while accumulation of MYb71 explains its successful colonization of worms. We next asked ourselves, whether the competitve advantage in worms would allow MYb71 to outcompete MYb11 in worms when following a biphasic life cycle that was iterated many times.

### Rapid environmental growth can overwhelm host selection

To evaluate whether host or environmental selection drive microbiota composition, we first simulated and then experimentally implemented repeated iterations of the biphasic life cycle. Abstracting the biphasic life cycle in a discrete mathematical model inspired by Lotka-Volterra dynamics, we focused on the two key selective pressures in the host and the environment: persistence in worms and competitive growth on agar (Fig. 3A; see equations 1.1 – 3.2 in the Methods section).

**Figure 3:**
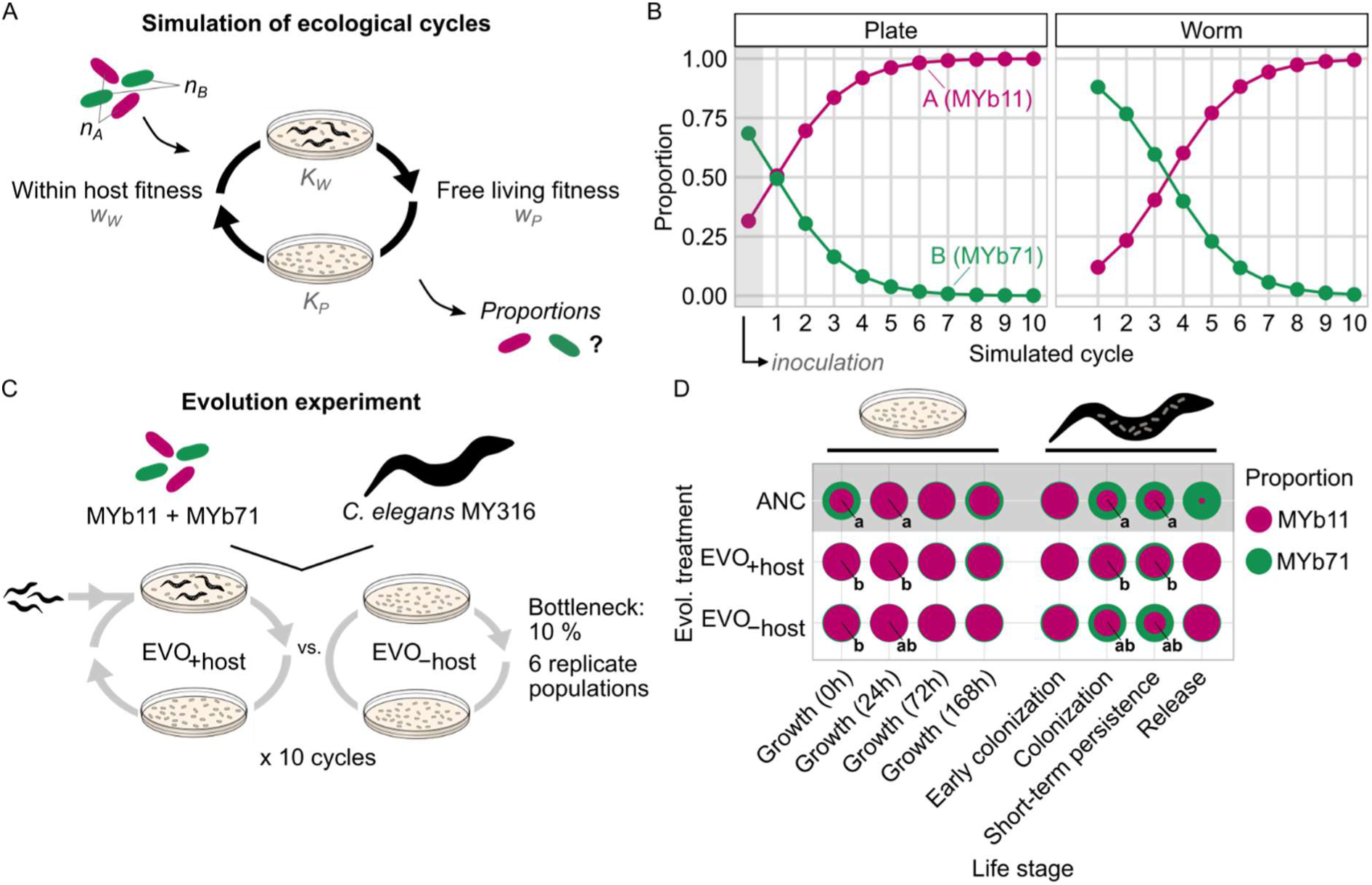
Repeated iterations of a biphasic life cycle lead to enrichment and host adaptation of the better environmental competitor MYb11. **A** A mathematical model of a simplified symbiotic life cycle was used to predict species proportions within the microbiota over time. Competitive fitness within the host or free-living environment and the carrying capacities of both habitats were estimated from empirical data (see Fig. 2). **B** Predicted species proportions after 10 iterations of the simulated symbiotic life cycle free-living (plate) and during host-association (worm). **C** In an evolution experiment, the two-member community of MYb11 and MYb71 was passaged either in a symbiotic biphasic life cycle with *Ce_*MY316 as a host (EVO+host) or as a no host control living on nematode growth agar. After 10 cycles, equivalent to 10 worm generations, the composition of ancestral and evolved microbiota communities was compared along the stages of the symbiotic life cycle. **D** Species proportions in ancestral, EVO+host and EVO–host communities in the different stages of the symbiotic life cycle are shown as nested circles. The area of inner (magenta) and outer (green) circles show the median proportion of MYb11 and MYb71, respectively (5 ≤ n ≤ 6). Statistical differences in species proportions between the ancestral and evolved communities are indicated by letters and calculated using a Generalized linear model with a binomial distribution and Tukey post-hoc tests.

Specifically, we simulated a two-member community initially seeded with both species, as in our ecological experiments (Methods; Supplementary Table S5). Bacteria from this community could then enter and persist in worms up to a maximum carrying capacity (𝐾_W_). Species abundances were weighted according to the species’ relative fitness, which we inferred from our experiments (𝑤_W_ ; Methods; Supplementary Table S5). This worm-associated community then seeded the environment in the next step, again according to species-specific free-living fitness (𝑤_P_) and a plate-specific carrying capacity (𝐾_P_). Simulating ten iterations of the life cycle, we observed that MYb11 rapidly increased in proportion on the agar plate and nearly reached saturation after about six cycles (Fig. 3B). Unexpectedly, we observed similar, yet lagging dynamics in simulated worms. Here too, MYb11 increased over time and after four simulated cycles it even dominated and subsequently approached fixation in the community (Fig. 3B). We hypothesized that the numeric advantage that MYb11 gains in the large environmental pool of symbionts, could influence species proportions in the host-associated community. To empirically test this, we used experimental evolution of the two-member community in a biphasic life cycle with a host-associated phase (EVO+host; Fig. 3C). In addition, we included a no host control (EVO-host) to compare the importance of environmental and host selection.

After serially passaging six independent communities of MYb11 and MYb71 in a life cycle with or without a host, we characterized relative fitness along the key phases of our biphasic life cycle. As predicted, MYb11 dominated communities grown on agar, at early time points even more so than ancestrally (Fig. 3D; Extended Data Fig. 3; Supplementary Table S6). During the different stages of host-association, too, community composition shifted towards greater proportions of MYb11. In light of the spill over from plates to worms that we had observed in our simulations, we expected that the growth advantage of MYb11 should lead to a particularly strong enrichment under evolution in the no host control. Intriguingly, however, MYb11 was significantly more enriched in worms (during colonization and persistence) after evolution in the host-associated treatment group (Fig. 3D; Extended Data Fig. 3; Supplementary Table S6). We therefore checked whether MYb11’s evolved advantage in worms came at a cost to MYb71 by comparing species abundances within ancestral and evolved communities. This showed that MYb71 abundances remained stable during colonization, if not slightly increased during short-term persistence, in evolved compared to ancestral communities (Extended Data Fig. 4; Supplementary Table S7). Therefore, we concluded that MYb11’s improved host-association did not result from improved competition, but rather evolutionary adaptation to the host.

Together, the combination of simulation and evolution experiments revealed that a MYb11’s growth advantage in the plate environment translates into an enrichment within hosts over ecological time and that this advantage can be enhanced by genetic adaption over evolutionary time. Finally, we aimed to assess whether evolved changes in microbiota composition had any influence on *C. elegans* hosts.

### Community evolution influences microbiota impact on host fitness

Improved bacterial fitness could be reached by either increasing abundance within hosts or by improving host, and indirectly bacterial, fitness. We therefore examined whether *Ce*_MY316 population sizes differed when worms were maintained on ancestral vs. evolved two-member communities (Fig. 4A). As a reference, we compared worm F1 population sizes reached on either MYb11, MYb71 or community lawns. Although not statistically different, population size reached on the co-culture resembled that of MYb71 more closely than that of MYb11 (Fig. 4B; Supplementary Table S8). Next, we tested whether evolution of a biphasic life history by the microbiota would change the impact on *C. elegans* hosts. Notably, worms maintained on evolved co-coltures had slightly greater population sizes than on ancestral bacteria (Fig. 4C, Supplementary Table S8). Consistent with worms reaching the highest population sizes in the presence of MYb11 (Fig. 4B), we found that the MYb11 enriched evolved microbiotas improve host fitness, in particular in those from the no host control (Fig. 4C). It therefore seems that shifts in microbiota composition during eco-evolutionary adaptation to the biphasic life cycle indirectly influence host fitness.

**Figure 4:**
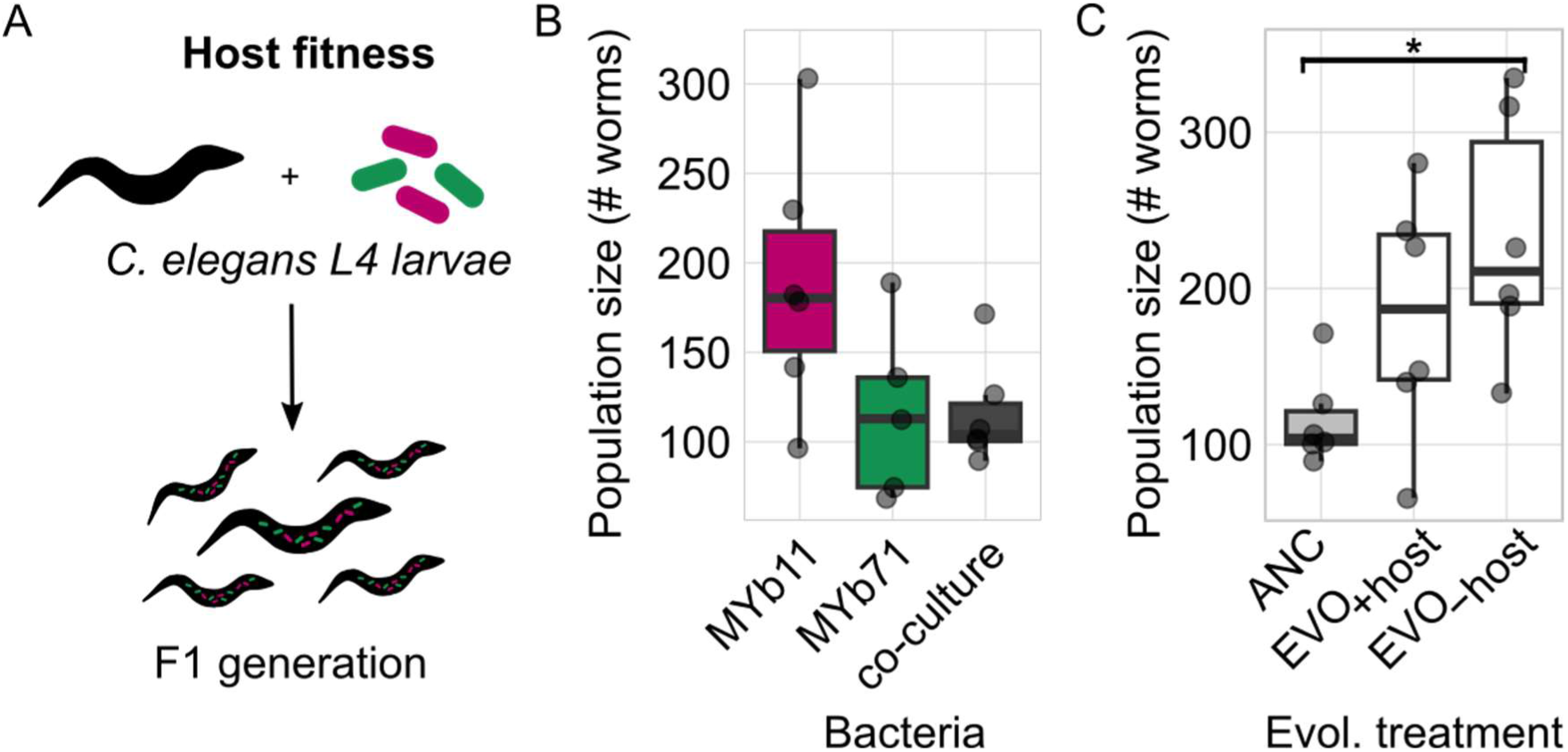
Microbiota composition and evolution affect host fitness. **A** Host fitness, i.e. worm population size, was quantified in association with mono- and co-cultures of MYb11 and MYb71. For this, the number of worms in the F1 generation produced by 5 *C. elegans* MY316 L4 stage within 3.5 days was counted. Replicate worm populations are shown as individual data points (5 ≤ n ≤ 6). Differences between **B** ancestral mono- and co-cultures of MYb11 and MYb71, as well as differences between **C** ancestral and evolved communities (EVO+host and EVO–host) were determined using ANOVAs with Dunnett post hoc comparisons (*p < 0.05).

## Discussion

In this study, we aimed to understand the importance of host and environmental selection on microbiota composition and function. To tackle this, we used a simple, two-member model community of MYb11 and MYb71 from the natural microbiota of *C. elegans* and performed competition experiments during environmental growth and worm colonization. Naturally, the nematode horizontally acquires a reproducible microbiota from its microbe-rich habitat, compost and rotting fruit^24, 25^. Although stochastic processes can affect worm colonization^26^, we here focus on two deterministic drivers of microbiota assembly: selection by host and environment and evolutionary adaptation. Our results show that over time, environmental selection can overwhelm host adaptation in this horizontally acquired microbiota.

Specifically, we found that competition within the environment is stronger than within the worm, although the relative fitness of the two microbiota members strongly depends on the time-point of sampling. Initially, MYb71 accumulates within worms thereby leading to an advantage in the host, whereas MYb11 shows an advantage in the environment due to its rapid growth. Experimentally imposing a biphasic life cycle with a host-associated phase in *Ce*_MY316 and a free-living phase on agar, we unexpectedly observed that the strong environmental competitor MYb11 dominates also the host-associated community over time. This is supported by a simple mathematical model and simulations of community composition over time. We therefore conclude that the numeric advantage of MYb11 in the environmental pool of symbionts overrides the fitness advantage of MYb71 within the host. Importantly, the resulting microbiota enrichment in MYb11 is reflected in host fitness, as population sizes of *Ce*_MY316 maintained on evolved communities come to resemble those of hosts on MYb11-only. Taken together, our findings suggest that environmental selection can dominate over host selection, and therefore plays a key role in shaping microbiota composition and function. In the following, we will first discuss how environmental selection overwhelms host selection. We then focus on the role of niche partitioning in maintaining species co-existence along a biphasic life cycle. Finally, we analyze how host adaptation of an environment-centric microbe enhances within host fitness.

### Environmental selection overwhelms host selection

It is commonly assumed that microbial taxa that are reliably found in association with a specific animal or plant are adapted to their host. Our findings, however, demonstrate that bacteria that are well-adapted to the host’s environment rather than the host itself can enrich within the host-associated community. Studying the microbiota isolates MYb11 and MYb71 revealed that the first is a better environmental competitor, whereas the latter can be considered a host specialist. In competition experiments along a biphasic life cycle, MYb11 outcompeted MYb71 on agar, while MYb71 gained a numeric advantage in the host by accumulating steadily over time. This is consistent with previous reports, where the *Ochrobactrum* isolate MYb71 is commonly found enriched in *C. elegans* microbiota communities of varying levels of complexity^12, 13, 17, 18^. While high insulin signaling in some *C. elegans* genotypes leads to a particularly pronounced enrichment of *Ochrobactrum*^21^, this taxon appears to be a more generally nematode-adapted. *Ochrobactrum* OTUs are not only found in *C. elegans*, but also as secondary symbionts of entomopathogenic nematodes ^27^ and as microbiota of predatory *Pristionchus* nematodes^28^. In contrast, the *Pseudomonas* isolate MYb11 was more competitive in the agar environment. While Pseudomonads are considered a core taxon of the natural *C. elegans* microbiota^17, 25^, they are comparatively rare both in natural and model *C. elegans* microbiota communities^12, 18^. Thus, we consider the isolate MYb11 an environmental specialist compared to MYb71. In spite of the fitness disadvantage during co-colonization, both bacterial species stably co-exist within the worm. This shows that distinct strategies may allow for maintenance during a biphasic life cycle, although they clearly differ in success.

While host- and environment-centric strategies thus coexist within our two-member community, we observed that environmental dominance can overwhelm host selection. This is in agreement with a recent theoretical model that inferred optimal fitness strategies for microbes that follow a biphasic life cycle^10^. This model suggested that a microbial population, which can grow within a host and a non-host environment and may transition between the two, may optimize fitness using two alternative strategies. Either, microbes should invest in increased replication in the environment, or decrease migration, and thus stay within the host. Our data suggests that the natural isolate MYb11 primarily follows the first strategy and reaps the benefits of rapid growth in the environment. As this bacterium increases in the symbiont source pool, is more likely to associate with the host, and transmission between hosts is thus increased. Importantly, an experimental study in zebrafish showed that dispersal of microbes between hosts can overwhelm selection by host immunity^29^. Thus, inter-host dynamics, including growth and survival in the free-living environment substantially affect within-host microbiota assembly.

### Evolutionary host adaptation of an environment-centric microbe enhances within-host fitness

Over evolutionary time scales, environment-centric bacteria can adapt to host-association within a microbiota. We observed that communities serially passaged in the presence of *C. elegans* hosts showed a greater enrichment in MYb11 than those in the no host control, i.e. maintained on agar alone. This suggested, that MYb11 increases fitness within hosts not only via environmental enrichment, but further via genetic adaptation after serial passaging.

In contrast to the ancestral environment-centric MYb11, the evolved MYb11 improved persistence in, and thus adaptation to, *Ce*_MY316 hosts. Strategically, we can link it to a theoretically expected optimal strategy of host-association, namely to decrease migration and stay with the host^10^. Our observations differ from that on *Lactobacillus plantarum* serially passaged in *Drosophila melanogaster*^30^. In this microbe-host pair, *L. plantarum* improved host-association via adaptation to the host diet, so rather by environmental enrichment. We hypothesize that MYb11 improved host adaptation as there was little scope for improved growth on the densely populated agar plates, while the opposite might apply to the probiotic *L. plantarum* adapting to *Drosophila*.

Taken together, we conclude that host adaptation of an environmental competitor, MYb11, not only provides an advantage in mono-association with the host but is robust in the context of a two-member community.

### Niche partitioning along the biphasic life cycle permits maintenance of species co-existence

Despite fluctuation along the biphasic life cycle, both species were maintained in our model community over ecological and evolutionary time scales. Although at different abundances, both MYb11 and MYb71 were consistently present in host and environmental communities. Co-existence of the two species was expected, as they have been co-isolated from a natural worm, have been co-cultured in *C. elegans* and on agar before, and can be maintained within a more complex microbiota context^12, 13, 18^. This is in line with the observation of prevalent bacterial co-existence, or positive co-occurrence, in the microbiota of *C. elegans*^17, 31^.

Classically, such patterns of co-occurrence are explained by niche partitioning as highly similar species should over time exclude each other by competition^32^. While our evolution experiment might have been too short to observe competitive exclusion, there are two reasons why niche partitioning along the biphasic life cycle can also explain the maintenance of both species. Firstly, we started with a host and an environment-centric microbe, so a system with already distinct favored niches. Thus, it is possible that both habitats can, at least temporarily, serve as a reservoir for either of the species. Secondly, we assume additional niche partitioning within the host. Similar to our observations here, the two main colonizers of the fresh water polyp *Hydra*, the bacteria *Duganella* and *Curvibacter* can stably co-exist on hosts, while *Duganella* outcompetes *Curvibacter in vitro*^33^. Distinct within-host niches might attenuate competition and thereby allow maintenance of MYb71 despite overwhelming MYb11 influx and host adaptation. Additional experiments would be required to uncover possible within-host niche differences between the two species and to explore to what extent they were emphasized and furthered character displacement, in our evolution experiment.

In addition to niche differentiation within the host and environment, we saw variation in microbiota composition over time, both in worms as well as on plate. As we started experiments with non-colonized worms, we may consider this temporal variation a succession series. Early on, MYb11 entered worms efficiently, yet as decreased in relative abundance as MYb71 accumulated over time. Despite an increasing disadvantage in the host, MYb11 was maintained in worms and even showed better persistence in the absence of bacterial influx (i.e. empty agar plates). This resembles observations from human gut microbiota successions in germ-free mice^34^. In mice, early-phase colonizers were maintained over time, although they were outcompeted by later-phase strains. In both systems, priority effects are thus expected to be weak, or of minor importance in structuring the community.

Variation in microbiota composition did, however, affect the metabolic functions of the microbiota^34^. Given the known differences in the functional repertoires of MYb11 and MYb71^14^, in our system, too, we can expect that differential abundance of two species should translate to different functional impact on the *C. elegans* host. Specifically, we expect that availability of thiamine and pantothenate, which only MYb11 but not MYb71 can produce^14^, should vary according to species abundances over time. While we did not track expression or activity of biosynthetic enzymes, we did observe that worms maintained on the evolved, MYb11-enriched communities, reached greater population sizes than on the ancestral, balanced community. Given that both vitamins are essential^35^, and that thiamine attenuated mitochondrial stress in a *C. elegans* disease model of dihydrolipoamide dehydrogenase deficiency^36^, we hypothesize that vitamin availability, could be driving the observed shifts in worm population growth.

Taken together, our observations on a simple, two-member community revealed that microbiota composition responds dynamically to a biphasic life cycle and that resulting species sorting feeds back on host fitness.

## Conclusion

Overall, our study showed how environmental selection can overwhelm host selection in the assembly of a two-member bacterial community associated with *C. elegans*. It highlights that eco-evolutionary dynamics in the symbiont source community directly feed back to the host-associated microbiota and can ultimately shape host fitness.

## Materials and Methods

### Bacterial and worm strains

In this study, we focused on a two-member microbiota community *Pseudomonas lurida* MYb11 and *Ochrobactrum vermis* MYb71, both key members of the natural *C. elegans* microbiota^12–14^. These species have been co-isolated from the natural *C. elegans* strain MY316 (*Ce*_MY316)^12^. Importantly, both bacteria can colonize worms and be grow on nematode growth medium agar (NGM)^37^, which allowed us to assay relative fitness (colony forming units/worm or CFU/agar plate) within hosts and the environment. For all experiments with ancestral bacterial isolates, we inoculated tryptic soy broth (TSB) overnight cultures (28°C, 150 rpm) with single bacterial colonies. We subsequently washed cells in M9 buffer and adjusted optical densities (OD; at 600nm) according to assay requirements. For competition experiments, bacterial species were co-inoculated at equal ODs. Relative species abundances were later determined via selective plating (on tryptic soy agar, TSA, with 10 µg/ml kanamycin), as only MYb71 is kanamycin-resistant.

The host in this study was *Ce*_MY316, recovered from a rotting apple in the Kiel Botanical Gardens^12^. For all experiments, we thawed *Ce*_MY316 from frozen stocks and performed colonization experiments with sterile and synchronized *Ce*_MY316. In preparation, *Ce*_MY316 was maintained on the non-colonizing food bacterium *Escherichia coli* OP50 according to standard protocols^37^.

### Bacterial growth

Bacterial fitness in the environment was measured in terms of population sizes after growth on NGM as (in colony forming units, CFUs). For this, we inoculated 6 cm NGM plates with 50 µl bacterial suspension at OD=0.1 and incubated plates at 20°C. CFUs were sampled, serially diluted, plated on TSA and counted at 24h, 72h and 168h.

### Bacterial colonization of worms

Bacterial fitness in the host was measured as CFU per worm. Briefly, five synchronized L4 stage *Ce*_MY316 were placed on a lawn of MYb11, MYb71 or both (6 cm NGM agar, 125 µl at OD600=2). After 3.5 days (20°C), worms were paralyzed and collected in M9 buffer with 0.025% Triton-100 and 25mM antihelminthic tetramisole. After washing on a custom-made filter tip washing system^23^, worms were collected in M9 buffer with Triton-100, and worm-free supernatant collected as background. Samples were then homogenized via bead beating and CFUs quantified via serial dilution and plating on TSA. CFU/worm were determined by subtracting background CFU from worm samples and dividing the total CFUs by the number of worms sampled.

### Early colonization, persistence, and release in worms

Early colonization, short- and long-term persistence, as well as release from the worm were measured as CFUs detected in or released from *Ce*_MY316 (see also^38^).

For early colonization, we placed L4 *Ce*_MY316 previously raised on *E. coli* OP50 on bacterial lawns (125 µl, OD600=2) for 1.5 hours. CFU/worm were then quantified as above (see bacterial colonization).

For short-term persistence and release, we raised synchronized L1 *Ce*_MY316 until L4 stage on bacterial lawns (125µl, OD600=2) for two days. For short-term persistence, we then washed worms on filter-tips and transferred to M9 buffer (300 µl). After taking a supernatant sample (100 µl), worms were then incubated for one hour (under shaking at 20°C) and a second supernatant sample (100 µl) was collected. Released bacteria were quantified as the difference between CFU in supernatant 1 and supernatant 2. CFUs maintained in worms were considered as having persisted in the short-term.

To assess long-term persistence, we transferred L4 *Ce*_MY316 previously raised on bacterial lawns of interest to empty NGM plates. After 24 hours of incubation (20°C), we quantified CFU/worm as above.

### Bacterial motility on agar

Colony expansion and swarming were quantified on NGM plates with either 0.5% or 3.4% agar. For both, 0.5 µl of bacterial suspension (OD600=1) were spotted on surface-dried agar plates. After 24 and 72 h, colony diameters were measured.

### Simulation model of microbiota composition during a biphasic life cycle

We built a mathematical model to assess how composition of a two-member community with MYb11 and MYb71 should respond to a biphasic life cycle over time. Mirroring our evolution experiment, we consider a community that experiences ten cycles of firsts associating with worm hosts and then growing on agar and use experimentally informed parameters accordingly (also see Figure 3A).

We use discrete Lotka-Volterra dynamics to describe bacterial abundances in hosts and on plate. For simplicity, we call MYb11 species A and MYb71 species B. We further assume that bacterial populations always reach the experimentally observed maximum population sizes (in L4 stage worms and on plates) and call these carrying capacities (on plate: 𝐾_P_, in worms: 𝐾_W_; Supplementary Table S6).

Preceding the first iteration of the life cycle, we assume that bacterial lawns are actively inoculated and at 𝐾_P_:

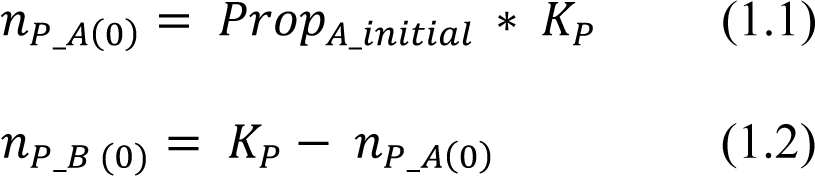

Species abundances on plate (𝑛_P_A_ and 𝑛_P_B_) at time zero are thus defined by the proportion of MYb11 in the experimental inoculum (𝑃𝑟𝑜𝑝_A_initial_) and 𝐾_P_. The remaining community members are considered to be MYb71.

For subsequent cycles, we allow bacteria from the plate environment to colonize worms according to their relative fitness in worms (𝑤_W_A_ and 𝑤_W_B_ = 1 − 𝑤_W_A_, respectively) and the carrying capacity of the host (𝐾_W_) in the host-associated phase:

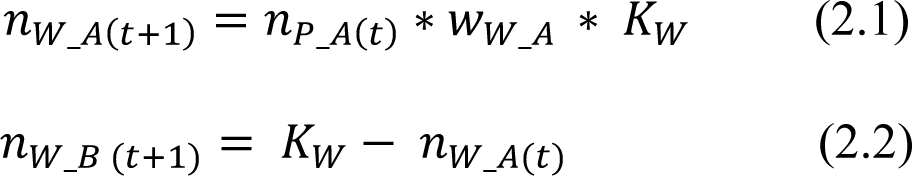

We thus consider that worms are maximally colonized at every iteration, as 𝐾_W_ is reached by default. This is in line with the observed consistent colonization of mono- and co-cultures observed experiments (see Fig. 1B). We then allow bacterial species to fill up worms, or plates, according to their abundance in the alternative habitat and their relative fitness.

In the free-living phase, bacteria grow in the environment according to their relative fitness on plates (𝑤_P_A_) and 𝐾_P_:

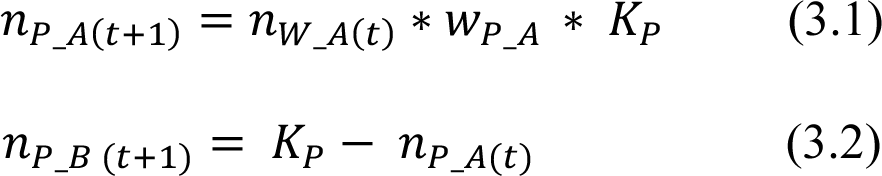

To analyse species proportions over time, we numerically simulated ten life cycles using the above equations in R. To eventually compare predictions with our evolution experiment, we inferred parameter values from empirical data collected from the ancestral bacteria (Fig. 2; Supplementary Table S6). Specifically, we assume that relative fitness in the host and thus transfer to the environment is a function of short-term persistence, as this most closely mirrors the transfer of host-associated bacteria sin the experimental evolution protocol. Thus, we estimate 𝑤_W_A_ and 𝑤_W_B_ from species proportions during short-term persistence in worms. In the evolution experiment, bacteria were then cultured on NGM for half a week.

We therefore assume that 𝑤_P_A_ and 𝑤_P_B_ are appropriately described by species proportions on agar after three days. Relative fitness thus always depends on the habitat-specific fitness and the influence of bacteria-bacteria interactions. As we focus on community dynamics in the absence of adaptation we assume that fitness values are static.

### Evolution experiment

Two-member communities of MYb11 and MYb71 were serially passaged along a biphasic life cycle on NGM either with *Ce*_MY316 (host treatment: EVO+host, 6 replicate communities) or without the host (no host control: EVO-host, 6 replicate communities) (Fig. 3C). Specifically, we used an established protocol for MYb11 evolution in mono-association with *Ce*_MY316^38^.

Briefly, NGM agar was seeded with mixed lawns of MYb11 and MYb71 (1:1 adjustment based on OD600) and cultured for 3.5 days (20°C). In the host treatment, ten L4 *Ce*_MY316 were placed on bacterial lawns and propagated until reaching the F1 generation (3.5 days). In no host controls, bacterial lawns were incubated without worms. We then collected bacteria from either worms or the agar environment at the end of every cycle. In the worm treatment, we transferred 10% of the community (bottleneck) to the next cycle and stored samples at -80°C. The no host controls were bottlenecked using a similar number of CFUs. We performed ten passages and subsequently analyzed the evolved communities.

Before phenotyping, we passaged thawed evolved communities along the respectively evolved life cycle and grew them on NGM agar as a common garden treatment to minimize bias introduced by freezing and thawing. Subsequently, cells were collected in M9 buffer and used in follow-up experiments (i.e. growth, colonization)

### Worm population growth

As a measure of host fitness on different bacterial lawns, we quantified worm population growth resulting from 5 L4 *Ce*_MY316 after 3.5 days. As in colonization assays, worms were maintained, washed and collected in M9 buffer. Worm samples were frozen in 48-well plates and photographed using a Leica dissecting scope (LEICA M205 FA). We counted worms in photographs using a semi-automated image analysis pipeline^38^.

### Statistical analysis

All statistical and mathematical analyses as well as data plotting were performed using R 3.6.1 in the RStudio environment^39, 40^ and ggplot2^41^. Normality and homogeneity of variances were checked by inspecting box plots of the data and, if absent, non-parametric tests were applied.

To assess fitness differences in the host and the agar environment, we compared CFU/worm and CFU/plate between MYb11 and MYb71 with those of the community using ANOVAs followed by Dunnett post-hoc tests that corrected for multiple comparisons by false discovery rate (FDR)^42^. We tested for differences in species abundances (log10(CFU)) during co-colonization and co-growth along the biphasic life cycle using a series of paired t-tests, again corrected by FDR. To infer differences in motility between the species and the two-member community using ANOVAs and Dunnett post-hoc tests (FDR-corrected). We tested for differences in species proportions along the biphasic life cycle by comparing ancestral to evolved (EVO+host and EVO-host) proportions using Generalized Linear Models with binomial distributions followed by Tukey post-hoc tests (FDR-corrected). To test for differences between species abundances log10(CFU) during colonization and short-term persistence of ancestral and evolved communities, we used Generalized Linear Models with Gamma distributions and Dunnett post-hoc tests (FDR-corrected). Finally, we compared worm population sizes on different bacterial lawns or ancestral vs. evolved communities using ANOVAs with Dunnett post-hoc comparisons (FDR-corrected).

## Data availability

All data is accessible alongside custom code via https://github.com/nobeng.

## Code availability

Custom code is available via https://github.com/nobeng.

### Acknowledgements

Acknowledgements: We thank F. Bansept, J. Zimmermann, P. Deines, and the Schulenburg lab for critical feedback, A. Czerwinski for lab support, the Kiel BiMo/LMB for access to their core facilities. Funding was provided by the Deutsche Forschungsgemeinschaft (DFG, German Research Foundation), Project-ID 261376515 – SFB 1182, Project A4 (NO, HS), the International Max-Planck Research School for Evolutionary Biology (NO), and the Max-Planck Society (Fellowship to HS).

## Author contributions

N.O.: conceptualization, methodology, investigation, software, data curation, formal analysis, visualization, writing - original draft, writing - review & editing. H.S.: conceptualization, supervision, writing - review & editing

## Additional information

Supplementary Information is available for this paper. Correspondence and requests for materials should be addressed to Nancy Obeng or Hinrich Schulenburg.

## Extended data figures

**Extended Data Fig. 1.**
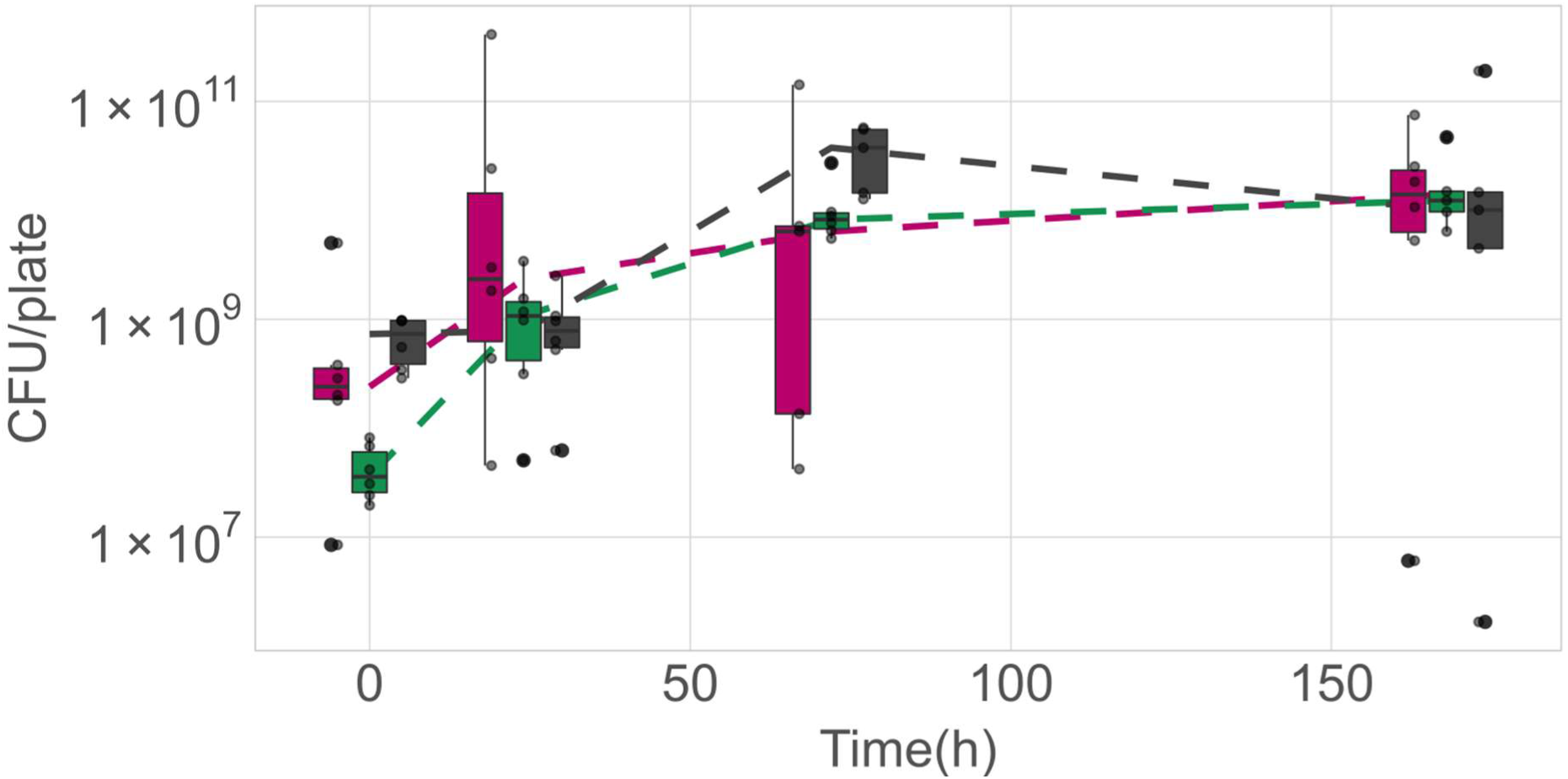
Growth of *Pseudomonas lurida* MYb11, *Ochrobactrum vermis* MYb71 and a co-culture on nematode growth agar over time. CFUs from replicate bacterial lawns (MYb11: magenta, MYb71: green, co-culture: grey) are shown as individual data points (5 ≤ n ≤ 6). Box plots summarize replicate cultures and show median (center line), upper and lower quartiles (box limits) and the interquartile range (whiskers). Dashed lines connect median CFUs per culture treatment over time.

**Extended Data Figure 2.**
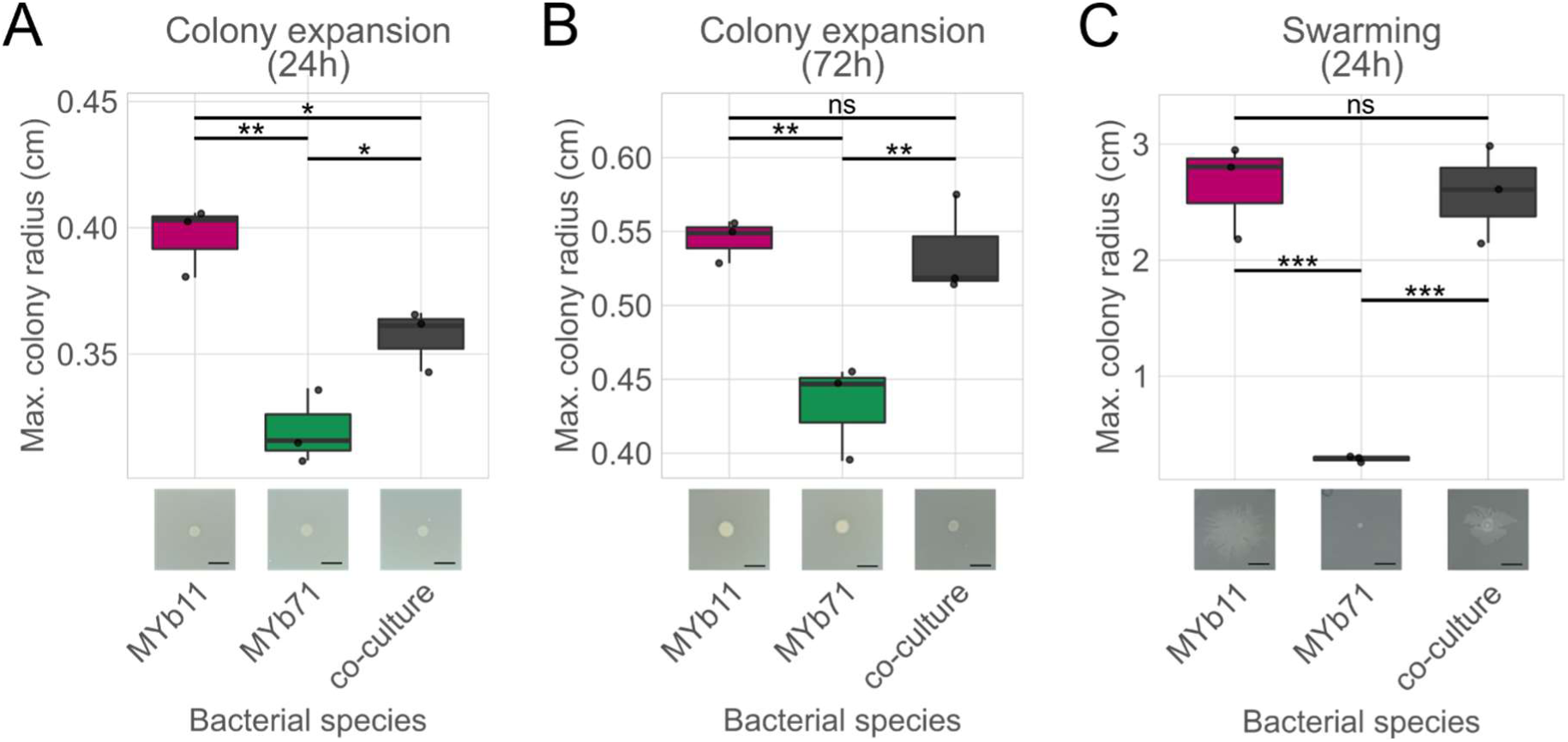
Motility of *Pseudomonas lurida* MYb11, *Ochrobactrum vermis* MYb71 and a co-culture on nematode growth agar. **A, B** Colony expansion on 3.4% agar was measured as colony radius after 24h and, B, 72 hours after spotting. **C** Swarming motility was assayed on 0.5% agar and max. colony diameter measured after 24h. Technical replicates are shown as individual data points. Species were compared using ANOVAs followed by Tukey post-hoc tests (***= P < 0.001, ** = P < 0.01, * = P < 0.05). Images of representative replicate colonies are shown along the x-axis, scale bars = 0.5 cm.

**Extended Data Figure 3.**
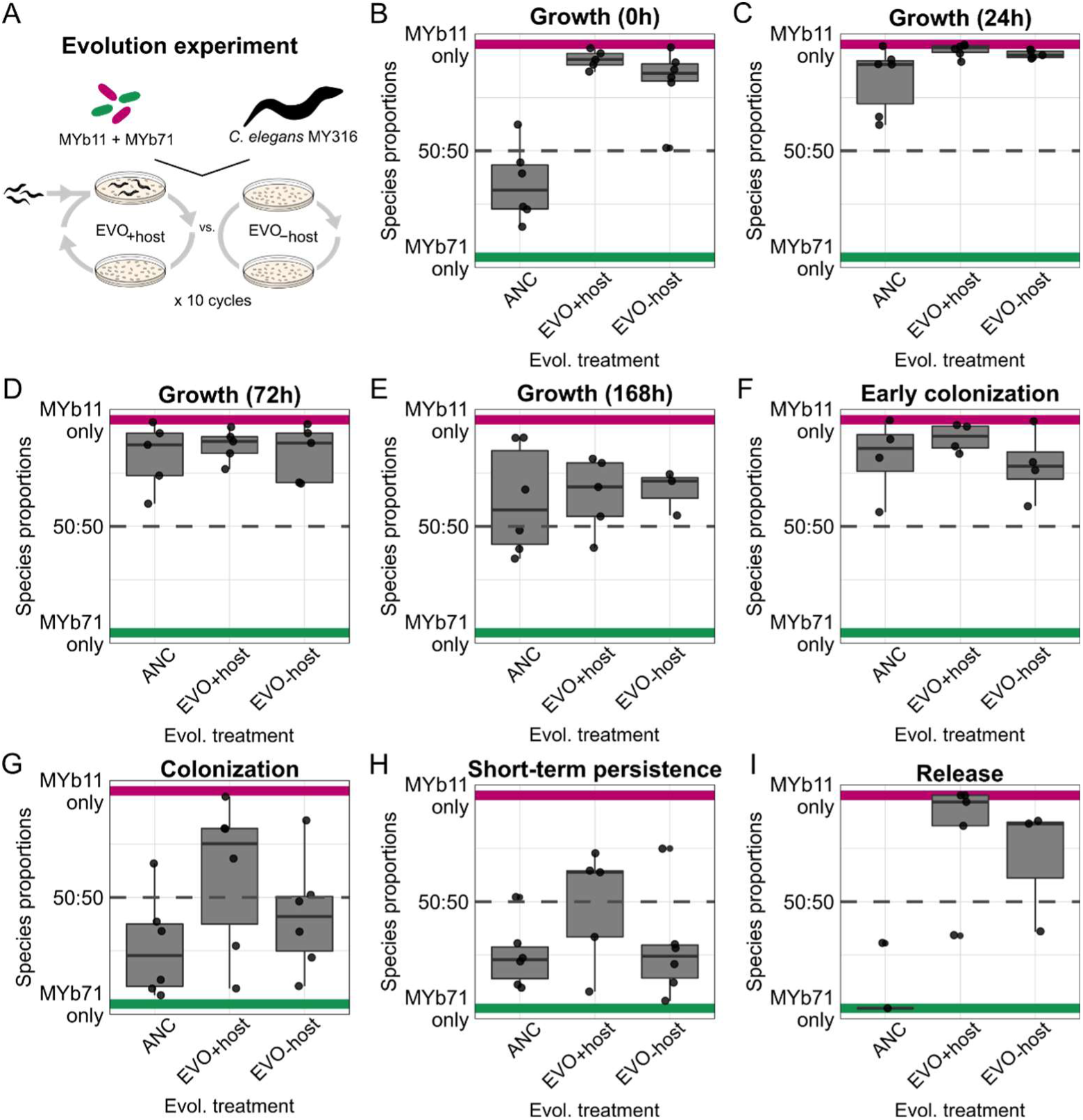
Proportions of *Pseudomonas lurida* MYb11 and *Ochrobactrum vermis* MYb71 in ancestral and evolved co-cultures across the stages of the biphasic life cycle. **A** Scheme of evolution experiment the analyzed communities were passaged in simplified from Fig. 1. Species proportions were quantified at, **B** inoculation of bacterial lawns (0h), **C** after growth on nematode growth agar for 24h, **D** 72h and **E** 168h. During host-association, species proportions were quantified during: **F** during early colonization of L4 stage Ce_MY316 (after 1.5h bacterial exposure), **G** established colonization of L4 stage *Ce*_MY316 (exposed to bacteria since L1), **H** short-term persistence in colonized L4 stage *Ce*_MY316 kept in buffer for 1h, and **I** release L4 stage *Ce*_MY316 into buffer within 1h. Replicate co-cultures are depicted as individual data points (5 ≤ n ≤ 6), dashed lines indicate 50:50 species proportions.

**Extended Data Figure 4.**
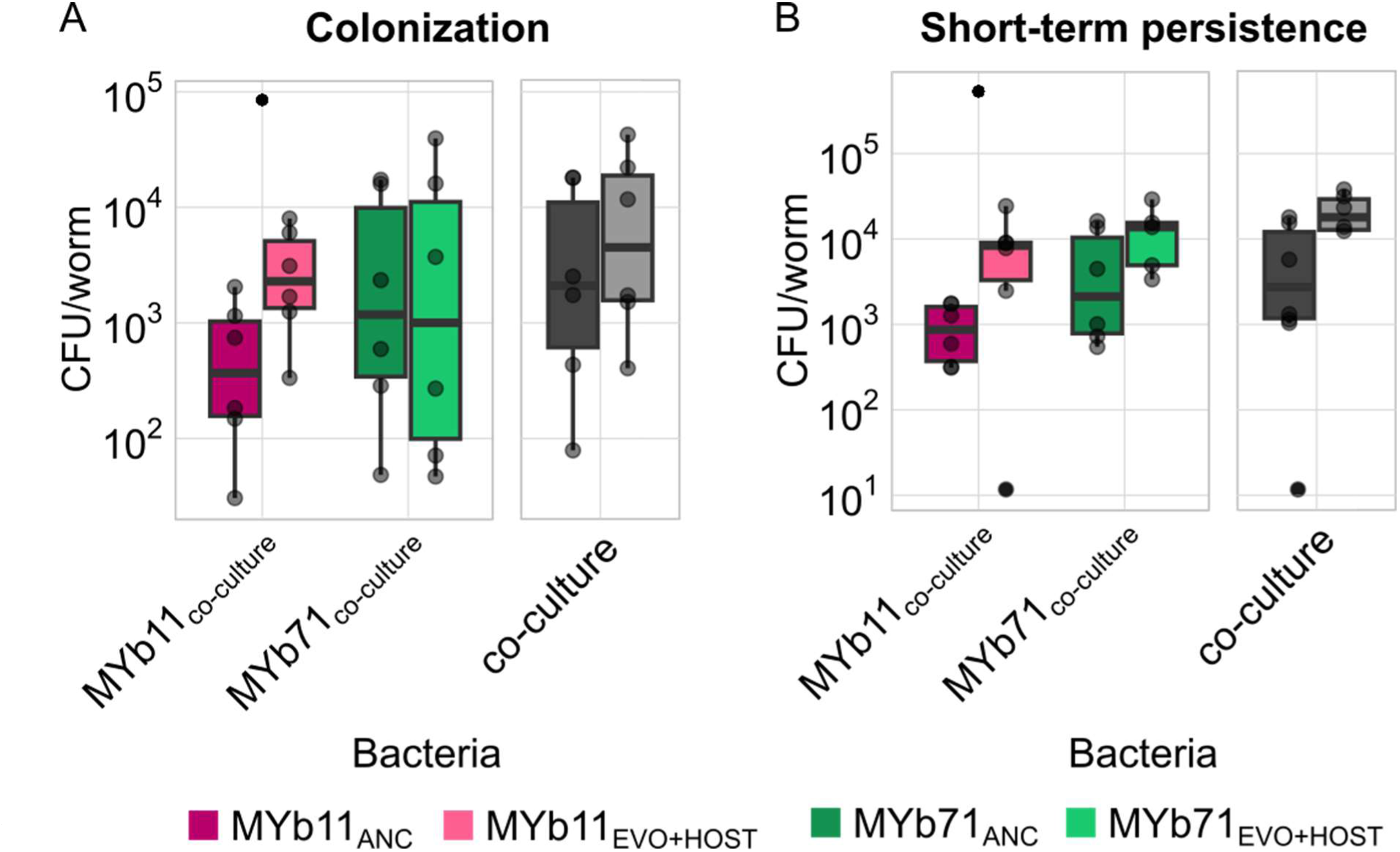
Microbiota composition during colonization and short-term persistence of ancestral and evolved co-cultures of *Pseudomonas lurida* MYb11 and *Ochrobactrum vermis* MYb71 in *Ce*_MY316. **A** Colonization, i.e. CFU/worm in L4 stage *Ce*_MY316 raised on an ancestral or host-adapted (EVO+host) co-cultures. **B** Short-term persistence, i.e. CFU maintained in colonized L4 *Ce*_MY316 that were kept in buffer for 1h. For both colonization and short-term persistence, MYb11 and MYb71 abundances within the co-culture are shown for the ancestral (dark magenta/green) and host-adapted (light magenta/green) communities (left box). In the right box, total colonization levels, i.e. the sum of MYb11 and MYb71 CFUs, are shown for ancestral and evolved bacteria. Replicates are shown as individual data points (5 ≤ n ≤ 6) and box plots summarize median (central line), upper/lower quartiles (box limits) and the interquartile range (whiskers).

## Supplementary tables

**Supplementary Table S1. Comparisons of MYb11 and MYb71 growth in monocultures with growth in co-culture using ANOVA followed by Dunnett post-hoc test.**

**Supplementary Table S2. Comparison of MYb11 and MYb71 colonization of Ce_MY316 L4 stage larvae in monoculture and in co-culture using ANOVA followed by Dunnett post-hoc test.**

**Supplementary Table S3. Comparison of relative abundances of MYb11 and MYb71 along the biphasic life cycle using paired t-tests. P-values shown are FDR-corrected.**

**Supplementary Table S4. Comparison of MYb11 and MYb71 motility (colony expansion and swarming) in monocultures and in co-culture using ANOVA followed by Tukey post-hoc tests.**

**Supplementary Table S5. Empirical parameters for simulation model of community composition along iterations of the biphasic life cycle.**

**Supplementary Table S6. Comparisons of species proportions in ancestral and evolved communities along the biphasic life cycle. For comparisons, Generalized Linear Models with Binomial distributions and Tukey post-hoc tests (FDR-corrected) were used.**

**Supplementary Table S7. Comparisons of species abundances of ancestral and evolved communities during colonization and short-term persistence. For this, GLMs with Gamma distributions as well as Dunnett post-hoc tests with FDR-correction were used.**

**Supplementary Table S8. Comparisons of worm population size on different species as well as ancestral vs. evolved community lawns. For this, ANOVAs with Dunnett post-hoc test and FDR correction were applied.**

